# Predicting mammal species suitable for Chagas disease spread

**DOI:** 10.1101/2025.02.13.638026

**Authors:** Maya Rocha-Ortega, Carezza Botto-Mahan, Gemma Rojo, Alex Córdoba-Aguilar

## Abstract

Given the risk of neglected diseases to humanity, can we predict how zoonotic diseases spread out? We have explored this by predicting mammal species reservoirs in the American continent that can act as potential blood-feeding sources of kissing bugs. Kissing bugs include 139 insect species of the Triatomine family that act as primary hosts of *Trypanosoma cruzi*, the causal agent of Chagas disease. *T. cruzi* also requires mammals as secondary hosts, yet our knowledge of which mammal species play such a role is uncertain. We analyzed trait data of mammal species documented as feeding sources to predict which other non-documented mammal species could then act as potential hosts. Using extreme gradient-boosted regression and having mammal traits as predictive variables, we found that 573 mammal species out of 1,923 species could be potential feeding sources of triatomines and, thus, reservoirs of *T. cruzi*. According to our analysis, triatomines overlap with these 573 mammal species on the basis of habitats, ambient temperature, and humidity. These insects also make use of mammals species that are heavier than 1 kg. Our estimate implies that controlling Chagas diseases is far more complex as the variety of potential mammal host species is extremely high. Also, the dynamics of Chagas disease may impose negative selection on many mammals in the wild.

The data can be gathered from: 10.6084/m9.figshare.28385720

## Introduction

Predicting and controlling zoonotic diseases are urgently needed tasks for humanity, as approximately 60–75% of emerging infectious diseases originate from zoonotic sources [1,2]. This wide realm of occurrence is partly explained by the fact that most animal pathogens can infect several host species, facilitating cross-species transmission and zoonotic spillovers [3–5]. However, the phenomenon of pathogen spillover relies upon a specific reservoir source that can host the pathogen at all times [4]. This particular reservoir connects to different target species or environments [6]. Moreover, we are still learning how zoonotic spillovers are influenced by interactions between reservoir hosts, pathogen exposure, and several factors that determine the susceptibility of the target host to infections, such as the life history and ecology of the interactant agents [7]. Although many humans, domestic animals, and wildlife diseases are assumed to be maintained in reservoir hosts, the implied reservoirs are rarely identified [6]. This information is essential to predict and control zoonotic spillovers.

Most emerging human diseases originate from non-human mammals [8,9], so we must predict which species are more involved during spillover events. Two predictors are ecological and/or life-history traits which influence mammalian species that harbor more pathogens [3]. More specifically, fast-living species—with small body sizes, short lifespans, rapid growth, and early reproduction—are expected to allocate relatively little energy to immune defenses against emerging pathogens [10], which grants them a higher reservoir competence [11]. Another factor favoring more pathogen encounters is the target host’s geographic range, as the wider the distribution, the greater the exposure to different pathogens [12]. One example of a typical target host is that of rodents given their traits associated with resilience to environmental disturbance: small-bodied, habitat generalists, rapid life histories, and extensive geographic distributions [13]. Finally, since certain traits are closely linked to ancestry (which explains why some taxonomic groups are more likely to harbor certain pathogens and thus serve as regular hosts), pathogen occurrence is linked to host phylogeny [14]. For instance, bats exhibit coevolutionary relationships with intracellular pathogens, granting them resilience to these infections and positioning them as reservoirs [15].

Over 10 million people are affected by Chagas disease [16]. This disease is caused by the protozoan *Trypanosoma cruzi*, which primarily moves between kissing bugs (Reduviidae, Triatominae) and terrestrial mammals [17]. Kissing bugs’ blood-feeding habit allows them to transfer *T. cruzi* parasites among domestic and sylvatic mammals [18]. The considerable diversity of triatomines includes species well adapted to environments such as urbanized and non-urbanized habitats [19].

Identification and monitoring of wildlife reservoirs of zoonotic diseases have been hampered for several reasons [20]. Consequently, we have an incomplete database of such reservoirs with a number of limitations: it stymies zoonosis prediction given by the biases of animal taxa linked to human disease; it obscures the evolutionary relatives of established zoonoses [21] as well as the interaction dynamics of vector-borne disease spread [22]. Despite these shortcomings, several recent models can still be used to identify and prioritize target reservoirs [21]: 1) network-based methods, which estimate a complete set of true host-pathogen interactions derived from a recorded network of associations; 2) trait-based methods that utilize observed relationships concerning host traits to identify species that conform to the morphological, ecological, or phylogenetic profiles of known host species associated with a given pathogen [20]; and, 3) genome sequences, which are particularly advantageous as they typically represent the initial (and often the only) data available for newly discovered viruses [21].

In this work, we have aimed to identify mammal species that may be exposed to and infected by *T. cruzi* in natural settings. We have used those species documented as part of triatomines’ feeding preferences to identify such predicted mammal hosts. We have also uncovered geographic areas where such predicted mammal hosts inhabit. Our work should serve as a guide to identify and predict susceptible reservoir mammals of *T. cruzi*.

## Material and Methods

### Chagas host status

Through a meta-analysis, we identified known mammalian hosts of Chagas disease, including both reservoir and target hosts, across distribution ranges in Mexico, Central America, and South America [17–19,23–38]. The host status is categorized as a binary trait: species are classified as confirmed positive infected with Chagas (1) or no record of infection (0). We compiled data from searches in the Web of Science and Scopus databases, focusing on studies published between 1980 and 2020. The search criteria included “(Chagas* OR Trypanosoma*) AND (mammal) AND (Americ*) EXCLUDING (Afric*). The aforementioned terms were employed in searching titles, abstracts, or keywords. The selection process for scientific papers was conducted in four distinct steps: 1) Identification: We conducted searches for scientific papers within the Web of Science (WoS), utilizing the specified keywords; 2) First screening: Abstracts were reviewed to select scientific articles that examined molecular testing of trypanosomiasis in mammalian species across the Americas or a review paper with confirmed cases of Chagas infection in mammals throughout America. During this stage, duplicate papers were also eliminated; and, 3) Final selection: Papers suitable for our analyses were determined. From 271 retrieved articles, we excluded 251 papers from our dataset after evaluating the titles and abstracts. This left us with 20 articles that examined mammalian communities as reservoirs for Chagas disease. For each paper, we sought to gather the following information: 1) authors; 2) country; and 3) taxonomic scope. Therefore, our analysis utilizing confirmed positive and non-positive record species data allowed us to identify species with a high likelihood of serving as hosts. Accordingly, we ended up with 83 known zoonotic Chagas species in the Americas (Supplementary Material Table 1). Then, we established the geographic range of triatomine distribution in the Americas based on triatomine records [39] and computed the concave polygon utilizing the *concaveman* function in R as a measure of similarity to the potential distribution of Chagas disease. Lastly, we compiled a list of 1,840 mammalian species that fall within the geographic range of Chagas disease for non-positive records sourced from GBIF mammal species.

### Sampling bias

We analyzed our dataset to uncover potential biases in representing host mammalian species. Such bias may emerge from an unequal sampling effort among mammal species, influenced by intrinsic biological traits, geographic distribution, taxonomy, and convenience. We gathered the number of Chagas-related publications for each mammal species to reflect the variation in research efforts across the mammal community. The number of publications was sourced from the Web of Science database through the execution of a query that incorporated the scientific binomial of the host species along with the search criteria “(Chagas* OR Trypanosoma*) AND (Americ*) EXCLUDING (Afric*)”.

### Traits linked with Chagas

For each of the 1,923 species in our dataset, a comprehensive set of life-history traits potentially correlating with carrier status was acquired from COMBINE [40] and EltonTraits [41]. Moreover, we employed IUCN range maps to ascertain the geographic range sizes (GRS) of native species in the Americas. In contrast, GRS was calculated on a global scale for introduced species. Furthermore, we incorporated factors related to climate [40,42], land cover type [43], and the global human modification index [44] within the territorial range of each species. Next, we calculated the average values of two bioclimatic variables, representing the warmest temperature and driest precipitation quarters for the carrier species across America [42]. To quantify phylogenetic traits, we obtained pairwise divergence times (phylogenetic distances) from TimeTree.org, a comprehensive knowledge base that has compiled and synthesized species divergence times from over 3,000 peer-reviewed studies [45]. The dimensionality of the matrix was reduced by mapping species within ordination space through a principal coordinate analysis performed in base R, yielding three dimensions that account for over 95% of the variation. Subsequently, correlated variables (r > 5) were eliminated utilizing the *corrplot* package in R. Two bioclimatic variables, temperature and precipitation, were retained as they are frequently employed to delineate the fundamental climatic requirements for mammal species on a global scale [46]. Finally, we retained two biogeographical variables: the geographical range size of distribution and altitude breadth; four ecological variables, which include the percentage of distribution across arboreal, grassland, agricultural, and urban habitats; and eight life-history traits, namely adult body length, brain mass, dispersal capacity, activity cycle, litter size, and the proportion of invertebrates, vertebrates, or plants in the diet.

### Data analysis

We employed extreme gradient-boosted regression (XGBoost) utilizing the xgboost package in R to analyze mammalian trait data to predict the probability of a species acting as a host for Chagas disease. XGBoost is classified as a machine learning algorithm that constructs an ensemble of weak decision trees. This methodology formulates a more robust predictive model by iteratively learning from weak classifiers, which are subsequently amalgamated into a stronger classifier; this process is called boosting [47,48]. Moreover, the XGBoost algorithm adeptly manages significantly unbalanced datasets by ascribing weights to positive labels, thereby enhancing class separability. In addition, it incorporates a regularization parameter that reduces the likelihood of overfitting to a limited number of positive labels. The capability to manage non-random patterns of missing data and its efficacy with considerably unbalanced data render the XGBoost algorithm the optimal choice for our research, proving particularly advantageous for our analysis, especially given that our dataset consists of a relatively small quantity of known hosts in comparison to the wide variety of mammal species under examination [48].

### Model performance and hyperparameter tuning

The accuracy of our model was evaluated through the implementation of nested cross-validation. We employed nested cross-validation to rigorously and impartially estimate the generalization error of our model ensemble [48]. A recent study examining the prediction of Leishmania hosts demonstrates that nested cross-validation is an effective resampling strategy for acquiring performance estimates when dealing with limited datasets. This methodology promotes the concurrent training and selection of models within a cohesive resampling framework, minimizing the risk of overfitting [48]. Additionally, by clearly delineating the uncertainty associated with predicted hazard hotspots, we emphasize regions where the reliability of predictions is compromised and identify potential knowledge gaps [10]. Nested cross-validation includes an inner loop specifically for hyperparameter tuning. Each iteration divides the dataset into five outer folds on the training dataset, consequently stratifying across three inner-fold cross-validations by host status category. This methodology yields predictions and evaluates the model’s comprehensive performance. The calibration of hyperparameters is executed utilizing Bayesian optimization, which aims to enhance a parameter space specifically designed to alleviate overfitting. This approach entails a minimal training rate, a reduced sample ratio used within the trees, a diminished feature ratio incorporated within the trees, a restricted maximum tree depth, and increased regularization [48].

Furthermore, the training dataset of each outer fold was partitioned into three inner folds, each comprising a training subset (67% of the outer fold’s training dataset) and an evaluation subset (33% of the outer fold’s training dataset). A target shuffling analysis assessed whether our model identified spurious correlations within the dataset. The nested cross-validation analysis was reiterated, with the response variables (host status) being randomly selected for each iteration. The nested cross-validation analysis was conducted repeatedly, during which the response variables (host status) were randomly selected for each iteration. Subsequently, the mean area under the receiver operating characteristic curve (AUC) was computed from the target shuffling experiment. The AUC serves as a classification metric that assesses the probability that the model output for a randomly selected positive label (known host) will exceed that of a randomly chosen negative label (an animal with no documented history of zoonotic Chagas infection). The AUC value ranges from 0 to 1, with an AUC of 1 signifying perfect classification of all samples, whereas an AUC of less than 0.5 implies that the model’s performance is no better than random chance [10,48].

### Chagas host trait profiles

We first identified key features using SHAP (Shapley Additive Explanations) via the R package SHAPforxgboost to assess Chagas’ host trait profiles. Shapley scores indicate the average marginal impact of each feature on predictions when considering all possible feature combinations. SHAP assesses local feature impacts by analyzing how each feature affects predictions for every data point in the dataset. We developed the model using 70% of the data to assess uncertainty in feature impacts, followed by calculating Shapley scores for the remaining 30% of observations. This process was repeated one hundred times, with different data subsets used in each cycle. Finally, we defined the trait profiles using SHAP partial dependence plots, demonstrating the link between a feature’s contribution to model predictions and its respective value in the dataset [48].

### Chagas mammal host predictions

Following reference [48], we identified certain animals as previously unrecognized zoonotic hosts for Chagas disease based on the Shapley value classification criterion. The aggregate of the Shapley scores across various features for each species corresponds to the prediction for that species, normalized by the average model prediction across all species. An animal was classified as a newly predicted host if more than 95% of the Shapley scores exceeded zero; in other terms, an animal was designated as a new predicted host if the expected probability of being a host surpassed the average likelihood for that model iteration for more than 95% of the model iterations [48].

## Results

After accounting for target shuffling, our model performed moderately well, with an average out-of-sample AUC of 0.94 (95% CI: 0.93–0.944) and an in-sample AUC of 0.98 (95% CI: 0.98–0.98).

### Predicted mammal species

A combination of biogeography, life history, and phylogenetic covariates ranked among the most significant model features when estimating global feature contributions (Fig. 1). Five hundred and seventy-three mammals were identified as potential new hosts. Among these newly predicted hosts, the majority belonged to the orders Rodentia (296 species), Primates (104 species), Carnivora (51 species), Eulipotyphla (41 species), Cetartiodactyla (25 species), Lagomorpha (23 species), Cingulata (18 species), Didelphimorphia (6 species), and Pilosa (6 species). Accordingly, while the order Primates represented the order with the highest scores, Rodentia represented the order with the highest number of species (Fig. 2). The newly predicted hosts exhibiting the highest cumulative Shapley scores, and included *Cheracebus torquatus*, *Pithecia pithecia*, *Callicebus personatus*, *Alouatta discolor*, *Saimiri cassiquiarensi*s, *Chiropotes albinasus*, *Callithrix geoffroyi*, *Brachyteles hypoxanthus*, *Plecturocebus miltoni*, and *Pithecia hirsute* (Supplementary Material Table 2).

**Figure 1.**
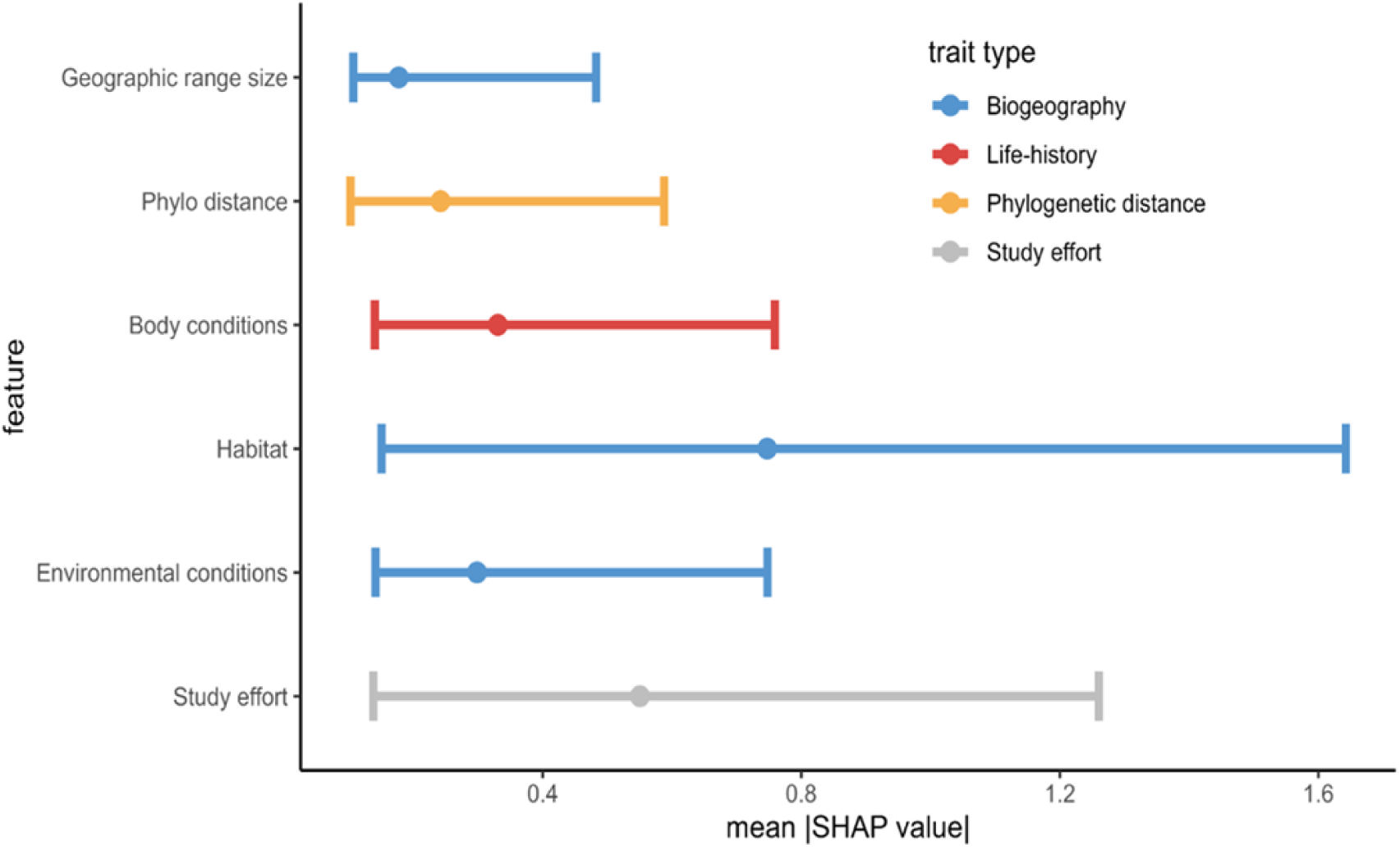
Traits linked to the probability of being a mammal host of triatomine’s feeding preferences. Biogeographical traits—represented in blue— include geographical range size (km²), the distribution percentage across arboreal, grassland, agricultural, and urban habitats, the warmest temperature, and the driest precipitation quarters. Life-history traits, shown in red, encompass adult body length and dispersal capacity. Phylogenetic distance is depicted in orange based on the location along PCoA ordination axes, while study effort is indicated in grey. Each point indicates the absolute value of the mean Shapley importance (mean |SHAP value|) for the trait across all mammals. The bars represent the absolute values of the 0.05–0.95 percentiles, and only features with 0.05 percentiles greater than zero are included.

**Figure 2.**
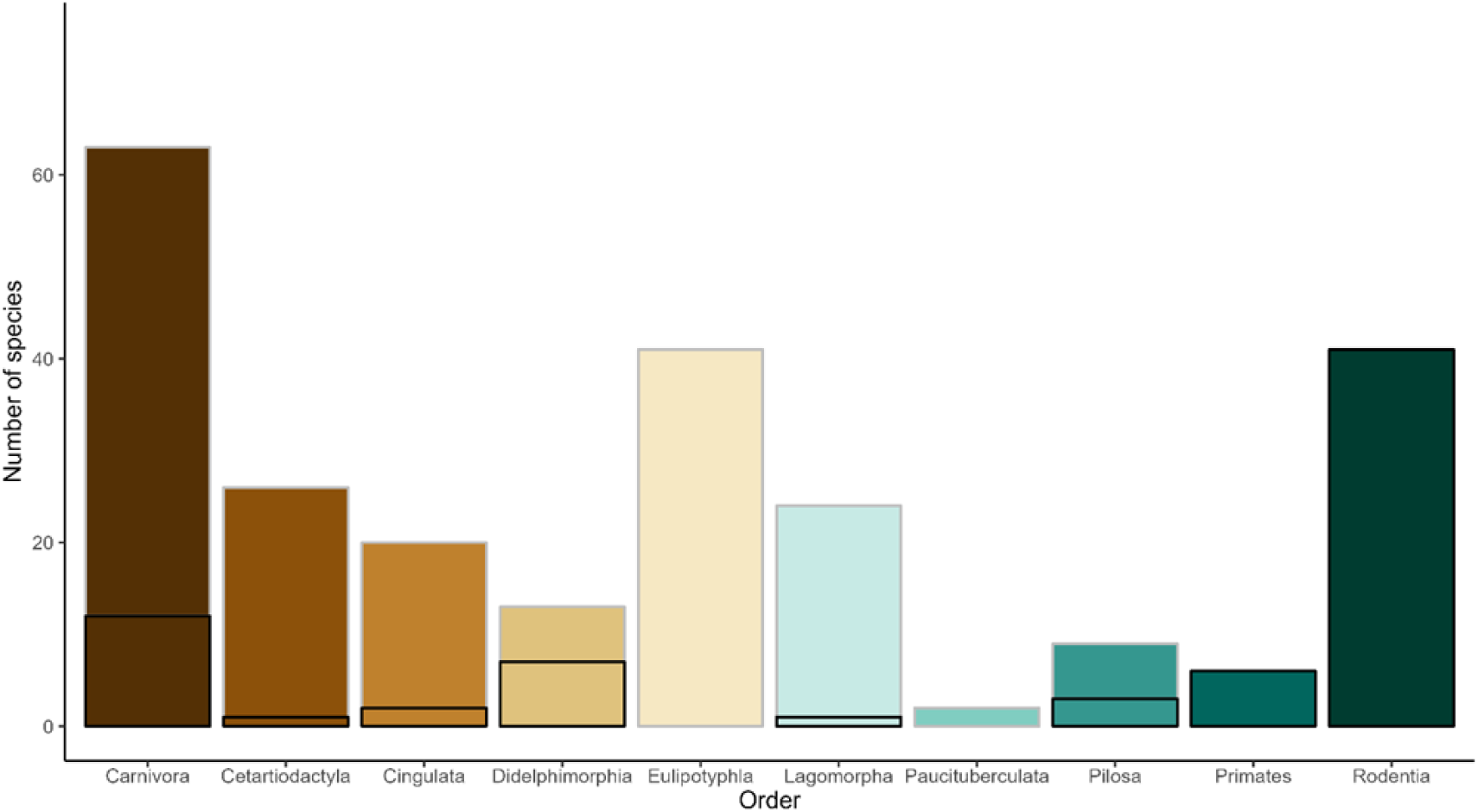
Taxonomic representation of mammalian species in our dataset. Black boxes represent the number of known mammal species hosts as a function of taxonomic order. Enlarged bars imply the number of new species predicted in our study.

### Mammal-associated traits as predictors of hosts

On average, habitat emerged as the most significant trait, followed by environmental and body conditions (Fig. 3). Thus, the warmest and driest locations, alongside crop and grassland cover areas, showed the highest cumulative Shapley scores. Medium-large-sized mammals weighing over 1 kg were identified as the species with the highest scores.

**Figure 3.**
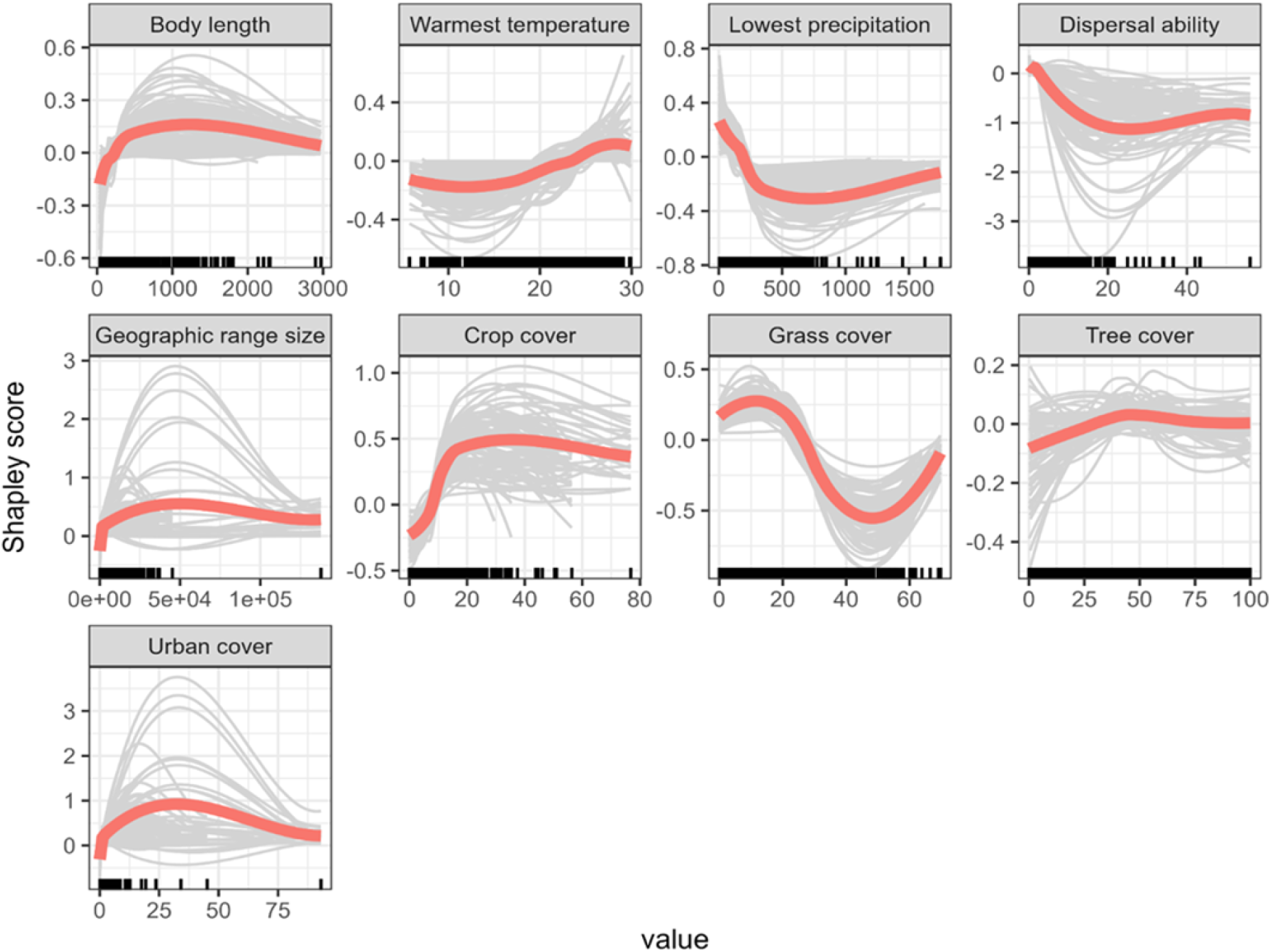
Shapley partial dependence plots illustrate the influence of each feature on *Trypanosoma cruzi* host’s status after considering the average effect of the other features in the model.

Characterized by geographical focal points, known host mammal species are primarily located in Mexico and Central America. In contrast to the known hosts, the geographical range of the newly predicted hosts predominantly concentrates in the Rocky Mountains, the Cordillera de Los Andes, Chaco, Mata Atlântica, and, mainly, the Amazon region (Fig. 4).

**Figure 4.**
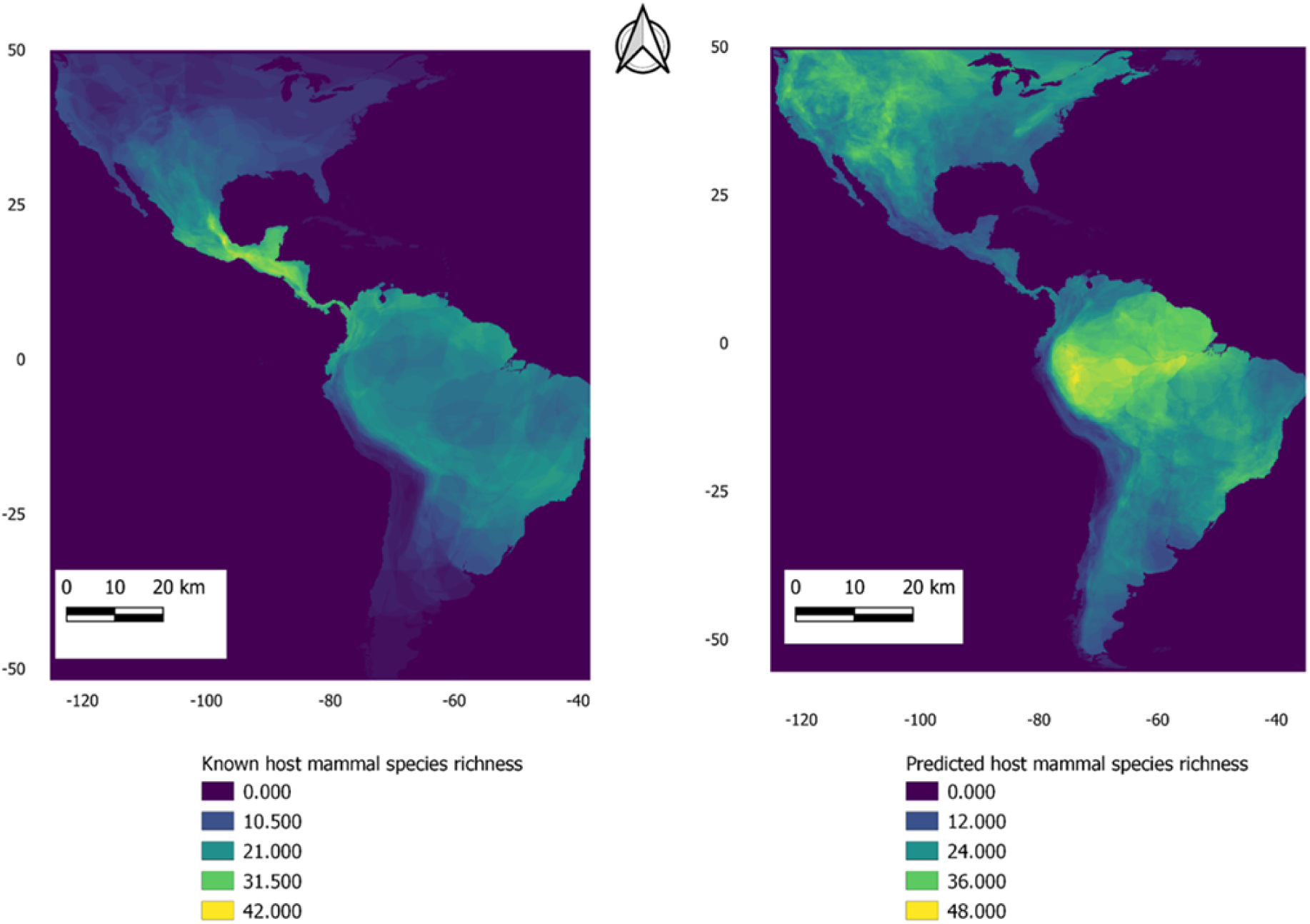
Distribution of zoonotic Chagas according to documented and predicted mammal hosts in the Americas.

## Discussion

Our study is the first to predict mammal species involved in the *T. cruzi*’s complex life cycle. The nearly six hundred mammal species is high considering the total number of mammals for the continent, which spans approximately 2200 species (although we used 1,923 species). This means that almost one-third of species on the continent can be suitable feeding sources for triatomines and, thus, potential hosts of *T. cruzi.* One first implication is that the ecology and control of Chagas disease are far more complex than thought, as the parasite has many host possibilities. One avenue of future research is to assess the dynamics of how triatomines and *T. cruzi* interact with mammals. One example of this exercise is the review of [31], who analyzed trypanosomatid infections across vertebrates in Chile. These authors were able to track which triatomine species participate in the transmission cycle of 25 mammal species according to the different Chilean regions. The geographical overlapping between the 25 mammal species and triatomines can then be used as a risk of Chagas disease. Another research avenue concerns the consequences of Chagas disease for wild mammals. Given that wild and domestic mammals can show signs of disease by *T. cruzi* infection [49], we hypothesize that *T. cruzi* may be a strong selective agent. The costs for mammals of *T. cruzi* infection may depend on the balancing forces between mammal resistance/tolerance and *T. cruzi* infectivity. We are unaware of the role of *T. cruzi* as a selective force for its reservoirs.

The geographical, habitat, environmental and body traits shared among the nearly six hundred mammal species can be explained by the ecological requirements of triatomines. As explained before, triatomines exhibit a wide repertoire of ecological requirements. For example, these insects exhibit thermal preferences, which can be found at particular altitude ranges. One case is *Triatoma dimidiata*, distributed in southern Mexico from 300 to 700 masl. Interestingly, at 700 masl, *T. dimidiata* is more likely to be infected with *T. cruzi*, whose isolate produces more acute symptoms in rodent subjects [50]. Habitat characteristics are also crucial for triatomines, and one case is that of human settlements. For example, triatomines preferred stacked brick and adobe enclosures as habitats in Arequipa, Peru [51], and palm trees in South America [52,53]. Environmental traits are possibly traits that have been more intensively studied with triatomines’ preferences (e.g. [54]). Ambient temperature and precipitation explain most triatomine species’ distributions (e.g. [55,56]). Triatomines prefer relatively high temperatures and dry environments (e.g. [57,58]). One would suppose that triatomines’ temperature and humidity preferences track those of the mammal species they use as feeding sources. Yet, the dietary niche breadth of triatomines does not explain their distribution range [55]. One interpretation of this is that the generalist nature of triatomines for host mammals makes the dietary requirements of triatomines less important compared to environmental conditions. As long as some particular temperature and humidity conditions are met, triatomines can use available feeding sources.

Finally, mammals that weigh more than 1 kg emerged as suitable hosts. Physiological, ecological and sampling factors can explain this. First, it has been demonstrated for other blood-feeding vectors, such as mosquitoes and ticks, that larger-bodied hosts are more attractive due to their higher CO_2_ emissions one of the main cues used to locate blood sources [59]. Triatomines also detect CO_2_ as a host-seeking cue [60], making it reasonable to assume that they prefer larger mammals, as these produce higher absolute volumes of CO_2_ [61]. Second, larger hosts have more stable body temperatures with larger skin surface areas, making feeding more accessible for triatomines [62]. Third, there may be a sampling effect as medium-sized mammals are typically the least studied compared to other body-size groups [63], which accounts for why many host species remain undetected.

We would like to discuss the size relation in more detail. Our research suggests that larger mammals are more likely carriers of Chagas disease, which impacts both fast- and slow-living species, much like viral infections [10]. The continuum of fast and slow life history traits outlines two pathways through which pathogen richness accumulates. This phenomenon may arise from the trade-offs between reproductive success and immune function, where 1) slow-lived species divert more energy towards immune responses, curtailing pathogen richness. 2) Increased exposure during extended survival and lifespan may enhance pathogen richness [64]. A host experiencing significant parasitism can achieve fitness by either (a) allocating resources towards immune resilience while prolonging reproductive efforts over a lengthened lifespan or (b) prioritizing rapid reproduction at the expense of immune investment, leading to heightened mortality and a reduced lifespan. Chronic infections like those may preferentially encourage greater immune investment and prolonged longevity[65]. Thus, medium-large mammal species may likely be subject to more pathogenic species throughout their lives, allocating resources to immunological resilience and allowing them to resist diseases such as Chagas.

We are unsure whether the habitat and environmental characteristics outlined are related to the probability of infection in triatomines. Specifically, we question whether certain habitat features and temperature and humidity ranges coincide with *T. cruzi* infections in these insects. This uncertainty arises partly because the infection status of triatomines (infected and non-infected) does not consider habitat and environmental variables. The most pertinent analysis in this context is the study conducted by [66], which demonstrated that the niche breadth of non-infected triatomines diverges from that of infected triatomines across seven species. This implies that infection in triatomines is not random, so some mammal populations may be more susceptible to infection if they coincide with infected triatomines that only occur under particular conditions. Consequently, future investigations should evaluate whether *T. cruzi* infects the anticipated mammal species delineated in this study.

In conclusion, our analysis is consistent with other research endeavors to predict zoonotic diseases by examining mammalian traits. For example, viruses related to zoonotic diseases are more prevalent in rodents and primates than in other mammalian species [67,68]. Furthermore, mammal species showed a fast-slow continuum of life histories, which, in another study, also harbored more virus zoonoses [10]. In alignment with these studies, our objective is to predict the potential for zoonotic diseases based on the host’s trait. Given the many possible mammalian hosts identified in our research, we advocate for implementing animal-targeted control measures for Chagas disease rather than approaches centered on human and triatomine populations [69]. In this regard, our list should serve as a guide to assess whether the predicted mammal species truly function as reservoirs and subsequently seek effective Chagas control strategies.

## Acknowledgements

This work was supported by grants from Fondo de Cooperación Chile México AGCID2024–Proyecto N°218 to AC-A, GR and CB-M; PAPIIT IN218824 to AC-A; ANID-FONDECYT-1221045 and ANID-ANILLO-ATE230025 to CB-M.

